# A glycoRNA switch for malignancy: SNORA73B activates TIAR-dependent oncogenic signaling in lung adenocarcinoma

**DOI:** 10.64898/2026.06.21.733650

**Authors:** Lulu Yang, Boyang Wang, Yufei Sheng, Zhuobin Deng, Jiali Liu, Zhiqi Hong, Lei Zheng, Chengwei Zhou, Wentao Hu, Zhaohui Gong

## Abstract

Although glycosylated small non-coding RNAs are emerging players in cancer, their functions in lung adenocarcinoma (LUAD) are largely unknown. We identify SNORA73B as a glycosylated small nucleolar RNA (glycol-snoRNA) that carries sialic acid-capped O-glycans in both normal lung epithelial and LUAD cells. SNORA73B is markedly elevated in LUAD, and its plasma levels distinguish early-stage LUAD from healthy controls with an area under the curve (AUC) of 0.7903. Subcellular fractionation reveals predominant nuclear localization. Functional assays demonstrate that SNORA73B depletion curbs LUAD cell proliferation, migration, and invasion, whereas its overexpression fosters these malignant phenotypes and accelerates tumor growth. Mechanistically, SNORA73B directly binds the T-cell-restricted intracellular antigen-related protein (TIAR), thereby enhancing TIAR protein abundance without affecting its mRNA levels. TIAR then recognizes the 3’-untranslated region (3’-UTR) of *MYC* mRNA to upregulate c-Myc, which subsequently augments AKT phosphorylation. Importantly, c-Myc knockdown largely rescues the oncogenic phenotypes and tumorigenesis induced by SNORA73B overexpression. Collectively, our data unveil a glycoRNA-dependent oncogenic axis SNORA73B-TIAR-c-Myc-AKT that drives LUAD progression. These findings position SNORA73B as a promising early diagnostic biomarker and a candidate therapeutic target in LUAD.

## Introduction

Lung cancer is the most common and deadliest malignancy worldwide^1^. Lung adenocarcinoma (LUAD) accounts for approximately half of all cases^2^. Despite advances in surgery and combination therapies, most patients are diagnosed at advanced stages, severely limiting treatment efficacy^3^. Identifying specific early diagnostic biomarkers is therefore an urgent clinical priority.

Non-coding RNAs (ncRNAs) are critical regulators of gene expression and disease progression^4,5^. Among them, small nucleolar RNAs (snoRNAs) canonically guide ribosomal RNA (rRNA) modification^6^ and also participate in DNA damage responses^7^ and protein interaction networks^8^. In cancer, snoRNAs exhibit dual roles: some act as tumor suppressors—for instance, SNORA13 enforces p53-dependent senescence by inhibiting ribosome biogenesis^9^—whereas others, such as SNORD17, promote hepatocellular carcinoma (HCC) through a NPM1/MYBBP1A positive feedback loop^10^. snoRNAs have been linked to lung cancer development and prognosis, underscoring their potential as both biomarkers and therapeutic targets^11^. However, systematic studies on snoRNAs specifically in LUAD remain scarce, and their mechanistic contributions are poorly defined.

Glycosylation, a fundamental post-translational modification, encompasses N-glycosylation, O-glycosylation, and other types^12^, covalently attaching glycans to proteins and lipids and profoundly influencing their functions. In cancer, glycosylation becomes widely dysregulated, yielding aberrant glycan structures that affect protein function^13^ and drive proliferation, invasion, metastasis, and immune evasion^14^. In lung cancer, glycosylation alterations arise across disease stages, correlating with invasion, metastasis and prognosis^15^, and stage-specific glycan profiles offer molecular signatures for early diagnosis and targeted intervention^16^. Yet research has focused almost exclusively on protein and lipid substrates, leaving the glycosylation of other macromolecules, particularly RNA, largely unexplored.

Traditionally, glycosylation was thought to be restricted to proteins and lipids; although post-transcriptional modifications increase RNA chemical diversity^17^, only a few transfer RNA (tRNA) monosaccharide modifications were identified^18,19^. Recent studies have challenged this view, showing that RNA can be glycosylated and that glycoRNAs participate in immune regulation^20,21^. Indeed, small ncRNAs, including Y RNA and snoRNAs, can carry N-glycans and be displayed on the cell surface, where they are recognized by sialic acid-binding immunoglobulin-like lectin (Siglec) receptors^22^. Cell-surface glycoRNAs further interact with P-selectin to modulate neutrophil-mediated inflammation^23,24^, and form complexes with heparan sulfate (HS) that directly bind and negatively regulate the pro-angiogenic factor VEGF-A165^25^. In breast cancer, surface glycoRNA levels inversely correlate with malignancy^21^, and serum glycoRNA signatures show strong tumour diagnostic potential^26^. Moreover, a glycoRNA-enriched neutrophil-like membrane delivery system has been developed for targeted therapy for abdominal aortic aneurysms^27^. GlycoRNA abundance also varies by cell type^28^, suggesting context-specific regulatory roles. Although initial efforts have mapped glycan attachment sites on glycoRNAs^29^ and predicted glycosylation motifs on small RNAs^30^, the systematic identification and functional characterization of glycosylated snoRNAs in LUAD have not been addressed.

Here, using metabolic labeling, small RNA microarrays, database analyses, and glycomics mass spectrometry, we identify and validate glycosylated SNORA73B (glyco-SNORA73B) in LUAD. Glyco-SNORA73B carries a high-molecular-weight O-glycan chain with terminal sialic acid modification and is significantly upregulated in LUAD cells, early-stage LUAD tissues, and plasma, highlighting its promise as an early diagnostic biomarker. Functional assays demonstrate that SNORA73B promotes LUAD cell proliferation, migration, invasion, and tumour growth in animal models. Mechanistically, SNORA73B drives malignant progression via the TIA-1-related protein (TIAR)/c-Myc/AKT signaling axis. Collectively, our findings establish glyco-SNORA73B as a novel candidate biomarker for early LUAD diagnosis and a potential therapeutic target.

## Results

### Glycosylated small RNAs are present in lung epithelial and LUAD cells

To detect glycoRNA, we metabolically labeled the LUAD cell line A549 and normal lung epithelial BEAS-2B cells with the sialic acid metabolic precursor Ac_4_ManNAz, which incorporates an azide handle into the sialic acid biosynthetic pathway. High-purity RNA was then subjected to click chemistry with a biotin-conjugated probe, and biotinylated glycoRNAs were visualized by denaturing gel electrophoresis and RNA blotting (**Fig. 1a**). After 36 h of Ac_4_ManNAz treatment, both cell types exhibited distinct steptavidin signals (Strep), whereas no signal was detected in the unlabeled controls (**Fig. 1b**). The hybridization signal was primarily distributed in the lower-molecular-weight region of the membrane, which, together with the SYBR-stained RNA profile (Sybr), indicated that glycoRNAs are preferentially enriched in the small RNA fraction. Notably, glycoRNA signals were observed in both normal and malignant lung epithelial cells, demonstrating that RNA sialylation is not restricted to cancer cells.

**Fig. 1.**
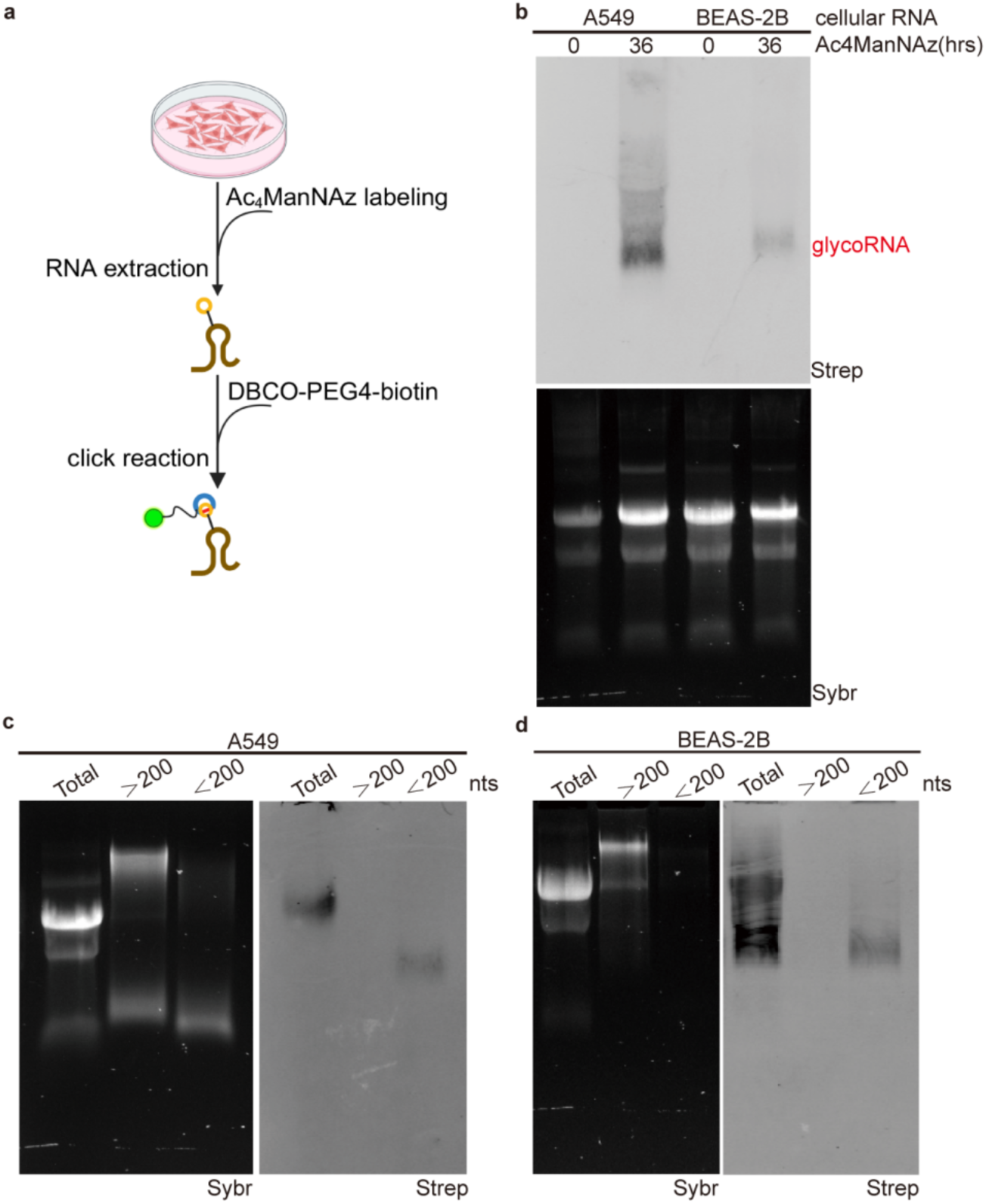
GlycoRNAs are enriched in small RNA fraction. (**a**) Schematic diagram of metabolic labeling. Cells were treated with the azide-containing sialic acid precursor Ac4ManNAz; following metabolic incorporation into sialylated RNA, extracted total RNA was biotinylated via click chemistry with DBCO-PEG4-biotin. (**b**) RNA blot analysis of total RNA from A549 and BEAS-2B cells with or without Ac4ManNAz treatment. Top: biotin signals using IRDye800 CW-labeled streptavidin (Strep). Bottom: SYBR Gold staining of total RNA prior to membrane transfer (Sybr). GlycoRNA signals are marked in red. (**c,d**) RNA blot detection of glycoRNAs in small (< 200 nt) and large (> 200 nt) RNA fractions obtained by silica column-based precipitation from Ac4ManNAz-treated A549 (**c**) and BEAS-2B (**d**) cells.

To resolve the size distribution of glycosylated RNAs, we fractionated total RNA into small (< 200 nt) and large (> 200 nt) populations using selective precipitation columns. GlycoRNA was detected exclusively in the small RNA fraction (**Fig. 1c,d**). We therefore conclude that glycosylated RNAs are predominantly species shorter than 200 nt.

### SNORA73B is a glycosylated and upregulated snoRNA in LUAD

To define the RNA species undergoing glycosylation, we performed small RNA microarray profiling on glycoRNA enriched by streptavidin pull-down from Ac_4_ManNAz-labelled A549 and BEAS-2B cells (**Fig. 2a**). Multiple small RNA classes—including microRNAs (miRNAs), tRNA-derived small RNAs (tsRNAs), precursor miRNAs (pre-miRNAs), tRNA and snoRNAs—were significantly enriched in the glycoRNA fractions (**Extended Data Fig. 1a,b**). Focusing specifically on snoRNAs, hierarchical clustering of the microarray data revealed a clear segregation between the snoRNA profiles of the Ac_4_ManNAz-enriched and input groups in both A549 and BEAS-2B cells, indicating specific snoRNA glycosylation (**Extended Data Fig. 1c,d**). We next identified glycosylated snoRNAs with |fold change (FC)| ≥ 2.0 and *P* < 0.05 upon Ac_4_ManNAz enrichment: 321 in A549 cells and 196 in BEAS-2B cells (**Fig. 2b**). Intersecting these two sets yielded 124 snoRNAs commonly glycosylated across both lines (**Fig. 2c**). To prioritize candidates with potential relevance to LUAD, we cross-referenced these 124 snoRNAs with 2,010 differentially expressed genes we previously identified in LUAD from TCGA data^31^. This analysis singled out SNORA73B as the only snoRNA meeting both criteria—glycosylation enrichment and differential expression in LUAD (**Fig. 2c**). Secondary structure prediction confirmed that human SNORA73B adopts the canonical H/ACA box architecture (**Fig. 2d**).

**Fig. 2.**
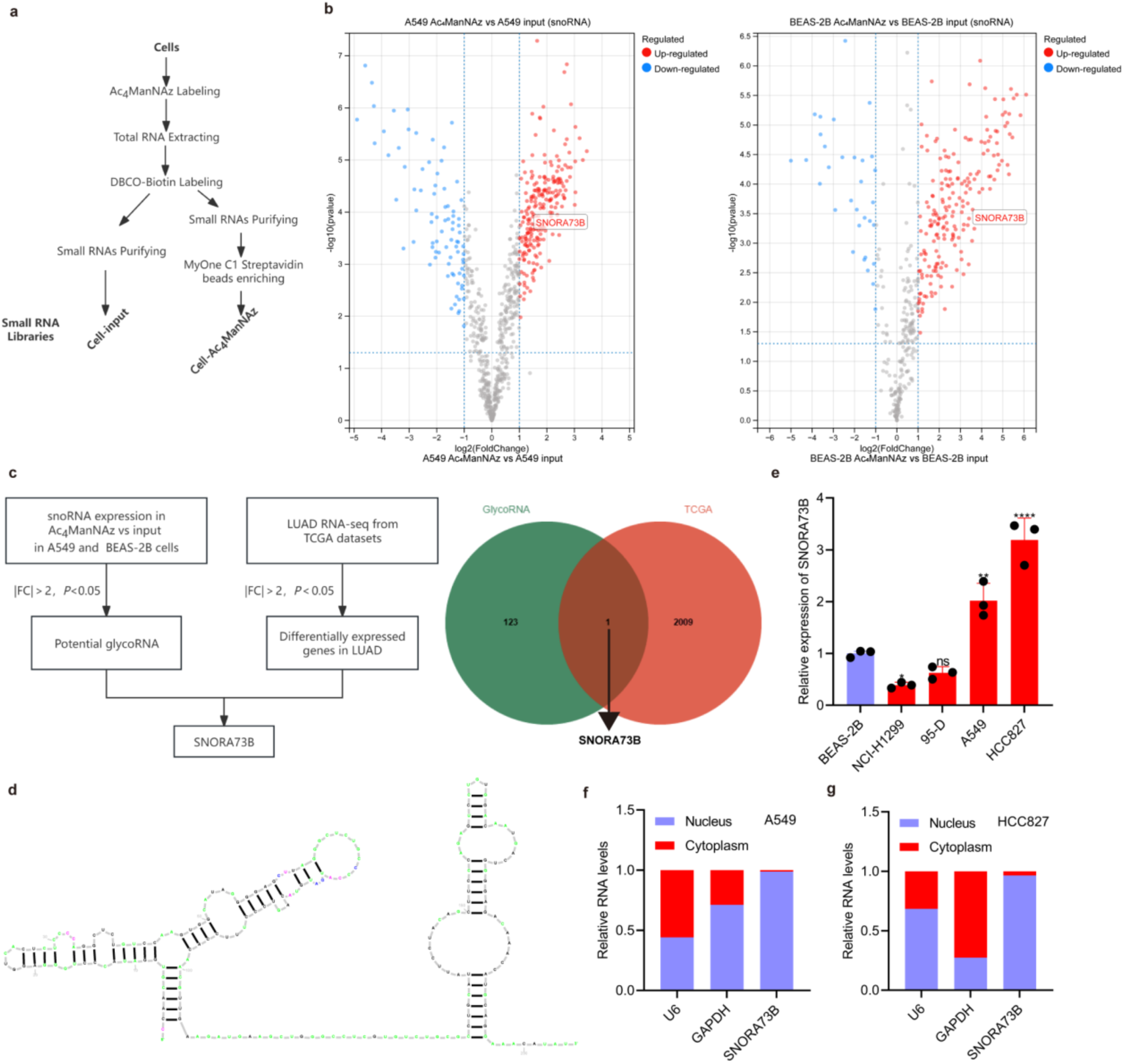
SNORA73B is a glycosylated snoRNA upregulated in LUAD. (**a**) Workflow for small RNA microarray analysis. Small RNA (< 200 nt) from input and streptavidin-enriched fractions of Ac₄ManNAz-labelled cells was profiled. (**b**) Volcano plots showing snoRNA enrichment in A549 cells and BEAS-2B cells comparing Ac₄ManNAz-enriched versus input fractions. Red and blue dots indicate significantly upregulated and downregulated glycosylated snoRNAs, respectively (thresholds: |FC| ≥ 2, *P* < 0.05). (**c**) Screening strategy and Venn diagram. The 124 glycosylated snoRNAs commonly upregulated in A549 and BEAS-2B cells with intersected with 2,010 differentially expressed genes from LUAD patients in the TCGA database (same thresholds as in **b**), identifying the SNORA73B as the target. (**d**) Predicted secondary structure of human SNORA73B (RNA Central database, https://rnacentral.org/). (**e**) RT-qPCR quantification of SNORA73B expression in normal lung epithelial cells (BEAS-2B) and LUAD cell lines (NCI-H1299, 95-D, A549, and HCC827). Data are presented as the mean ± standard deviation (SD), n = 3 independent experiments. Statistical significance was determined by one-way ANOVA. (**f,g**) Nuclear and cytoplasmic distribution of SNORA73B determined by RT-qPCR after subcellular fractionation in A549 (**f**) and HCC827 (**g**) cells. Ns, not significant, \**P* < 0.05, \*\**P* < 0.01, \*\*\*\**P* < 0.0001.

RT-qPCR analysis across four LUAD cell lines showed that SNORA73B was significantly upregulated in A549 and HCC827 cells relative to BEAS-2B normal lung epithelial cells (**Fig. 2e**). Nuclear-cytoplasmic fractionation demonstrated that SNORA73B localizes predominantly to the nucleus in both A549 and HCC827 cells (**Fig. 2f,g**), consistent with its predicted nucleolar residence of a box H/ACA snoRNA. Together, these results establish SNORA73B as a conserved, glycosylated snoRNA that is markedly overexpressed and primarily nuclear in LUAD cells, providing a basis for functional characterisation.

### SNORA73B bears high-molecular-weight sialylated O-glycans

The markedly retarded electrophoretic migration of glycoRNAs suggested modification by large glycan structures. To define the glycans present on SNORA73B, we performed a glycomic mass spectrometry on RNA from A549 cells expressing non-targeting (sh-Control) or SNORA73B-targeting shRNA (sh-SNORA73B-2) (**Fig. 3a**). Both O- and N-glycans were detected in both groups (**Fig. 3b,c**; **Extended Data Fig. 2a,b**), demonstrating the structural diversity of endogenously glycosylated RNAs. Further quantitative comparison revealed that sialylated O-glycan subtypes were significantly altered upon SNORA73B knockdown, indicating that depletion of SNORA73B specifically affects O-glycan sialylation. These data provide the first glycomic evidence that SNORA73B is modified with both N- and O-glycan structures.

**Fig. 3.**
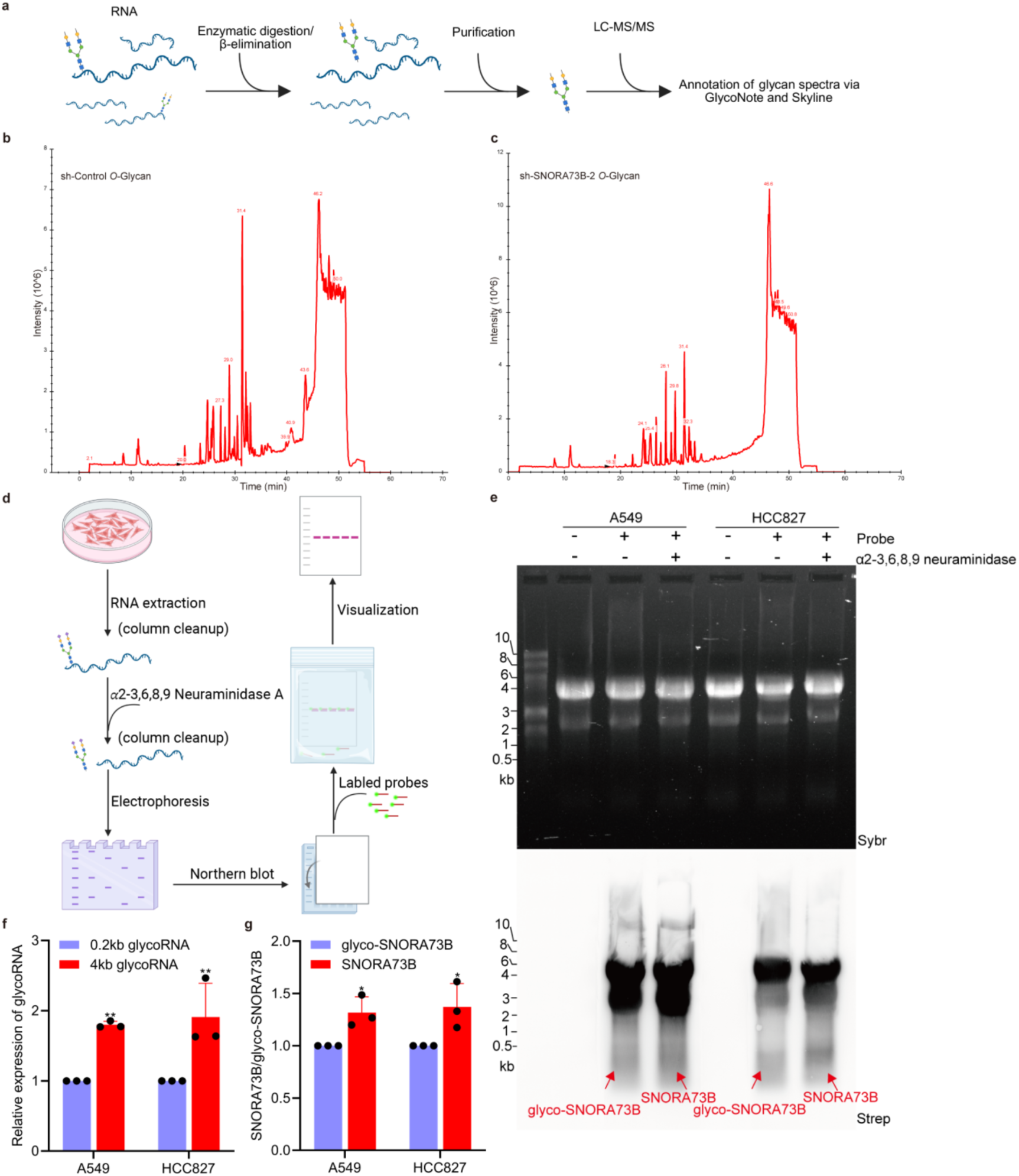
SNORA73B carries high-molecular-weight, terminally sialylated O-glycans. (**a**) Schematic of glycan release and purification for glycomic mass spectrometry analysis. (**b,c**) Total ion chromatograms (TICs) of O-glycans from sh-Control (**b**) and sh-SNORA73B-2 (**c**) samples. (**d**) Experimental strategy: total RNA was treated with α2-3,6,8,9 neuraminidase, and SNORA73B was detected by RNA blotting. (**e**) RNA blot of SNORA73B in A549 and HCC827 cells with or without sialidase treatment. Glyco-SNORA73B (high-molecular-weight) and deglycosylated SNORA73B (low-molecular-weight) are indicated in red. (**f**) Quantification of the large (∼4 kb) and small (∼0.2 kb) bands in the untreated samples shown in **e**. (**g**) Quantification of the deglycosylated SNORA73B band intensity in sialidase-treated versus untreated samples. Data are presented as the mean ± SD, n = 3 independent experiments. Statistical significance was determined by two-way ANOVA (**f,g**). \**P* < 0.05, \*\**P* < 0.01.

To determine the size and sialylation status of the O-glycan chains on SNORA73B, we combined broad-specificity sialidase digestion with RNA blotting (**Fig. 3d**). Total RNA from A549 cells was treated with α2-3,6,8,9-sialidase to remove terminal sialic acid residues, resolved by denaturing gel electrophoresis, and detected with a SNORA73B-specific probe. In the untreated samples, the SNORA73B signal concentrated in a high-molecular-weight region (∼4 kb), with only a faint signal at the low-molecular-weight region (∼200 nt) (**Fig. 3e,f**). Sialidase treatment caused a pronounced shift of the signal to the ∼200 nt position (**Fig. 3g**), demonstrating that removal of sialic acids increased electrophoretic mobility. We conclude that SNORA73B is modified with high-molecular-weight O-glycan chains that carry terminal sialic acids, rather than with small glycosyl groups, establishing a structural basis for its functional characterization.

### SNORA73B holds diagnostic potential for early-stage LUAD

To evaluate the clinical relevance of SNORA73B, we quantified its expression in paired tumour and adjacent normal tissues from 76 LUAD patients. SNORA73B was significantly upregulated in tumour tissues (**Fig. 4a**), and this elevation was already prominent in stage I disease (n = 66; **Fig. 4b**). We further measured plasma SNORA73B levels in plasma from 66 LUAD patients and 27 healthy controls (HCs). Plasma SNORA73B abundance was significantly higher in LUAD patients than in HCs (**Fig. 4c**) and was likewise elevated in stage I patients (n = 63; **Fig. 4d**). Thus, SNORA73B is overexpressed in both tissues and plasma from the early stages of LUAD.

**Fig. 4.**
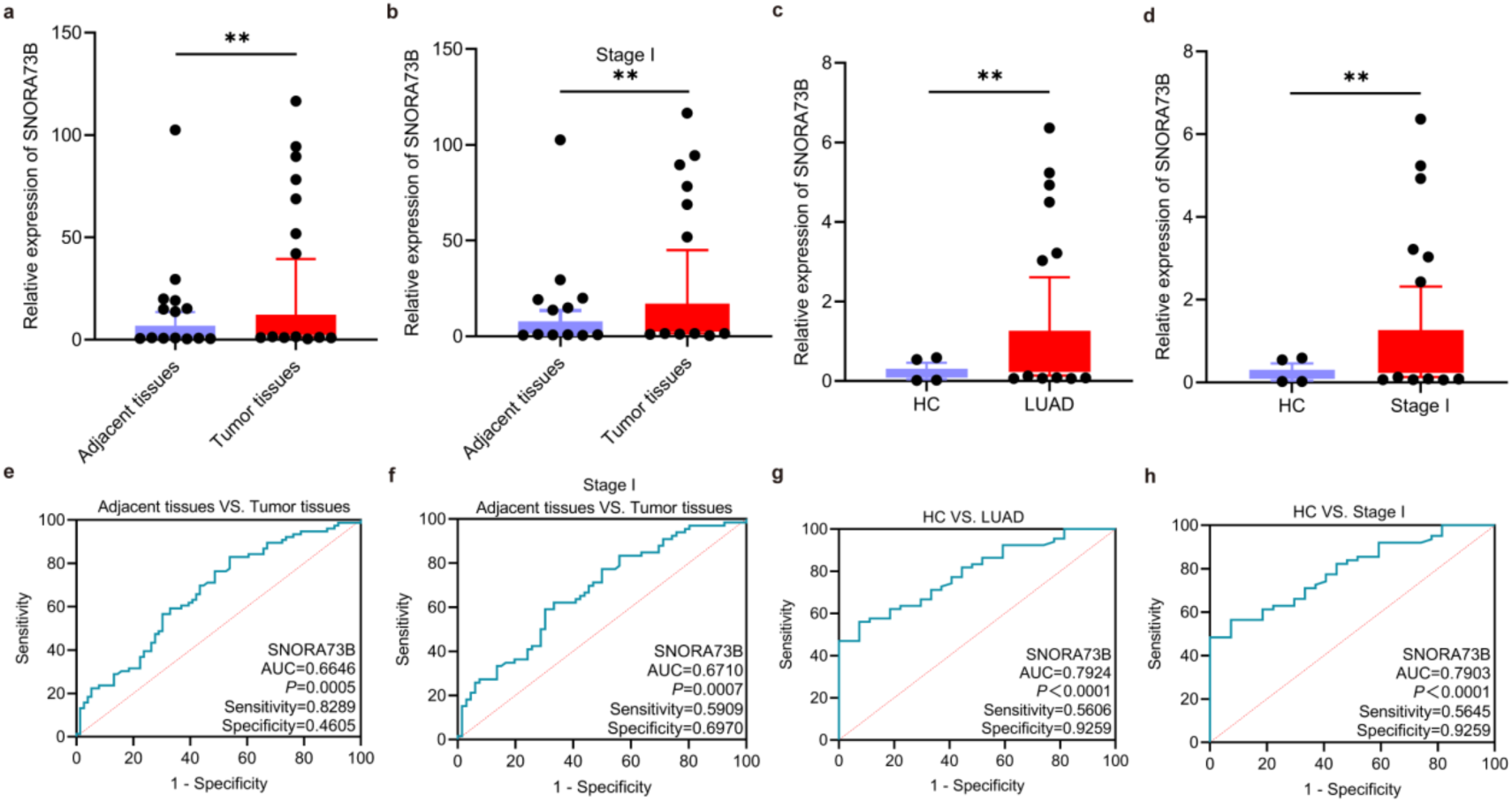
SNORA73B expression and diagnostic performance in LUAD tissues and plasma. (**a**) RT-qPCR analysis of SNORA73B in paired tumour and adjacent normal tissues from 76 LUAD patients. (**b**) Paired tissue comparison in 66 patients with TNM stage I LUAD. (**c**) Plasma SNORA73B levels in healthy controls (HCs, n = 27) and LUAD patients (n = 66). (**d**) Plasma SNORA73B levels in HCs (n = 27) and TNM stage I LUAD patients (n = 63). (**e,f**) ROC analysis of SNORA73B expression for discriminating LUAD (**e**) or TNM stage I LUAD (**f**) from normal tissues. (**g,h**) ROC curves based on plasma SNORA73B expression for discriminating LUAD patients (**g**) or TNM stage I LUAD patients (**h**) from HCs. Statistical significance was determined by two-tailed unpaired Student’s *t*-test(**a,b,c,d**). \*\**P* < 0.01.

Receiver operating characteristic (ROC) analysis was performed to assess diagnostic performance. In tissue, SNORA73B discriminated LUAD from normal with an area under the curve (AUC) of 0.6646 (sensitivity: 82.89%, specificity: 46.05%) and stage I tumors with an AUC of 0.6710 (sensitivity: 59.09%, specificity: 69.70%) (**Fig. 4e,f**). In plasma, the AUC was 0.7924 for distinguishing LUAD patients from HCs (sensitivity: 56.06%, specificity: 92.59%) and 0.7903 for stage I patients versus HCs, maintaining a specificity of 92.59% (**Fig. 4g,h**). These results indicate that SNORA73B possesses diagnostic efficacy in both tissue and plasma, with notably high plasma specificity for early-stage LUAD, supporting its potential as a non-invasive early diagnostic biomarker.

### SNORA73B promotes LUAD malignant progression

To investigate the function of SNORA73B in LUAD, we generated three shRNAs targeting SNORA73B and an overexpression (OE) construct. Transfection into A549 and HCC827 cells showed that sh-SNORA73B-2 achieved the greatest knockdown efficiency (**Fig. 5a**), and SNORA73B OE significantly upregulated SNORA73B expression (**Fig. 5b**). CCK-8 and colony formation assays demonstrated that SNORA73B knockdown markedly inhibited proliferation and clonogenicity, whereas SNORA73B overexpression promoted both in LUAD cells (**Fig. 5c,d** and **Extended Data Fig. 3a-d**). Moreover, Transwell assays revealed that SNORA73B knockdown impaired cell migration and invasion, while SNORA73B overexpression of significantly promoted them (**Fig. 5e,f** and **Extended Data Fig. 3e,f**). These data indicate that SNORA73B facilitates LUAD cell proliferation, migration and invasion.

**Fig. 5.**
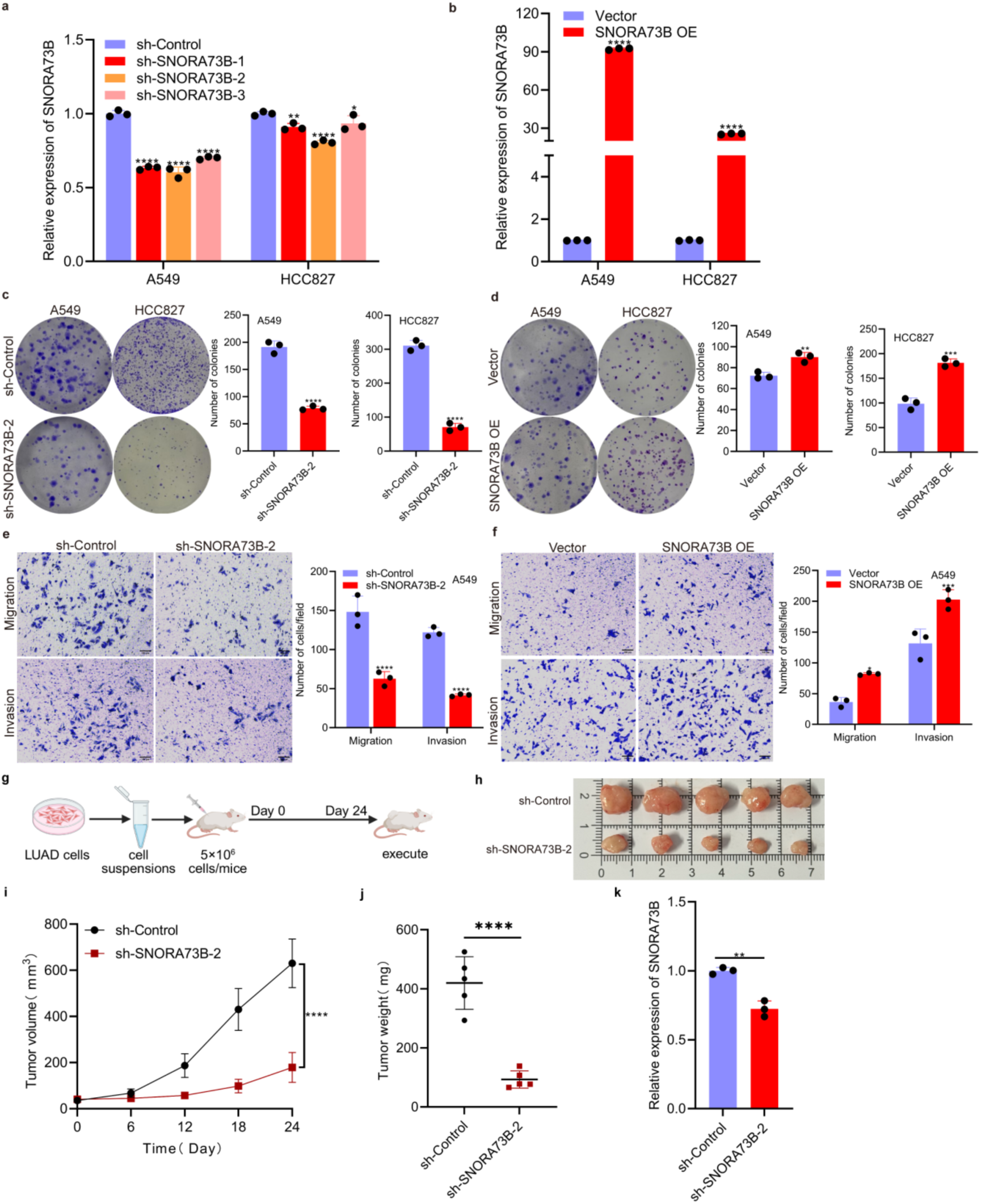
SNORA73B promotes malignant progression of LUAD. (**a,b**) RT-qPCR validation of SNORA73B expression in A549 and HCC827 cells transfected with three independent shRNAs or a non-targeting control (**a**) and with SNORA73B overexpression (OE) or empty vector (**b**). (**c,d**) Colony formation assays of A549 and HCC827 cells upon SNORA73B knockdown (**c**) or overexpression (**d**). (**e,f**) Transwell migration and invasion assays of A549 cells with SNORA73B knockdown (**e**) or overexpression (**f**). Scale bars, 100 μm. (**g**) Schematic diagram of the subcutaneous xenograft experiment. (**h**) Representative images of xenograft tumors excised from nude mice injected with sh-Control or sh-SNORA73B-2 cells (n = 5 mice per group). (**i**) Effect of SNORA73B knockdown on tumor volume over time. (**j**) Effect of SNORA73B knockdown on tumor weight at endpoint. (**k**) RT-qPCR analysis of SNORA73B expression in xenograft tumors. Data are presented as the mean ± SD, n = 3 independent experiments (**a-f,k**) or n = 5 mice per group (**h-j**). Statistical significance was determined by two-way ANOVA (**a,b,e,f,i**) or two-tailed unpaired Student’s *t*-test (**c,d,j,k**). \**P* < 0.05, \*\**P* < 0.01, \*\*\**P* < 0.001, \*\*\*\**P* < 0.0001.

To determine whether SNORA73B is required for tumour growth *in vivo*, we established subcutaneous xenografts in nude mice using A549 cells expressing sh-Control or sh-SNORA73B-2 (experimental scheme in **Fig. 5g**). The results showed that SNORA73B knockdown significantly reduced tumour volume and weight (**Fig. 5h-j**). Further RT-qPCR analysis of excised tumors confirmed lower SNORA73B expression, consistent with the diminished tumorigenicity (**Fig. 5k**). These results demonstrate that SNORA73B is essential for LUAD tumour formation *in vivo*.

### SNORA73B binds TIAR and upregulates its protein abundance

To identify SNORA73B-interacting proteins, we performed biotinylated RNA pull-down with A549 lysates coupled to mass spectrometry (**Fig. 6a**). The sense strand probe enriched 181 proteins relative to the antisense control (**Extended Data Fig. 4a,b**). Intersection of these candidates with three RNA bind protein (RBP) databases (RBPsuit, catRAPID, and ENCORI) yielded six proteins shared hits: TIAR (T-cell-restricted intracellular antigen-related protein), WT1-associated protein (WTAP), serine/arginine-rich splicing factor 9 (SRSF9), transformer 2 alpha homolog (TRA2A), nuclear RNA export factor 1 (NXF1), and ATP-binding cassette subfamily F member 1 (ABCF1) (**Fig. 6b and Supplementary Table 1**).

**Fig. 6.**
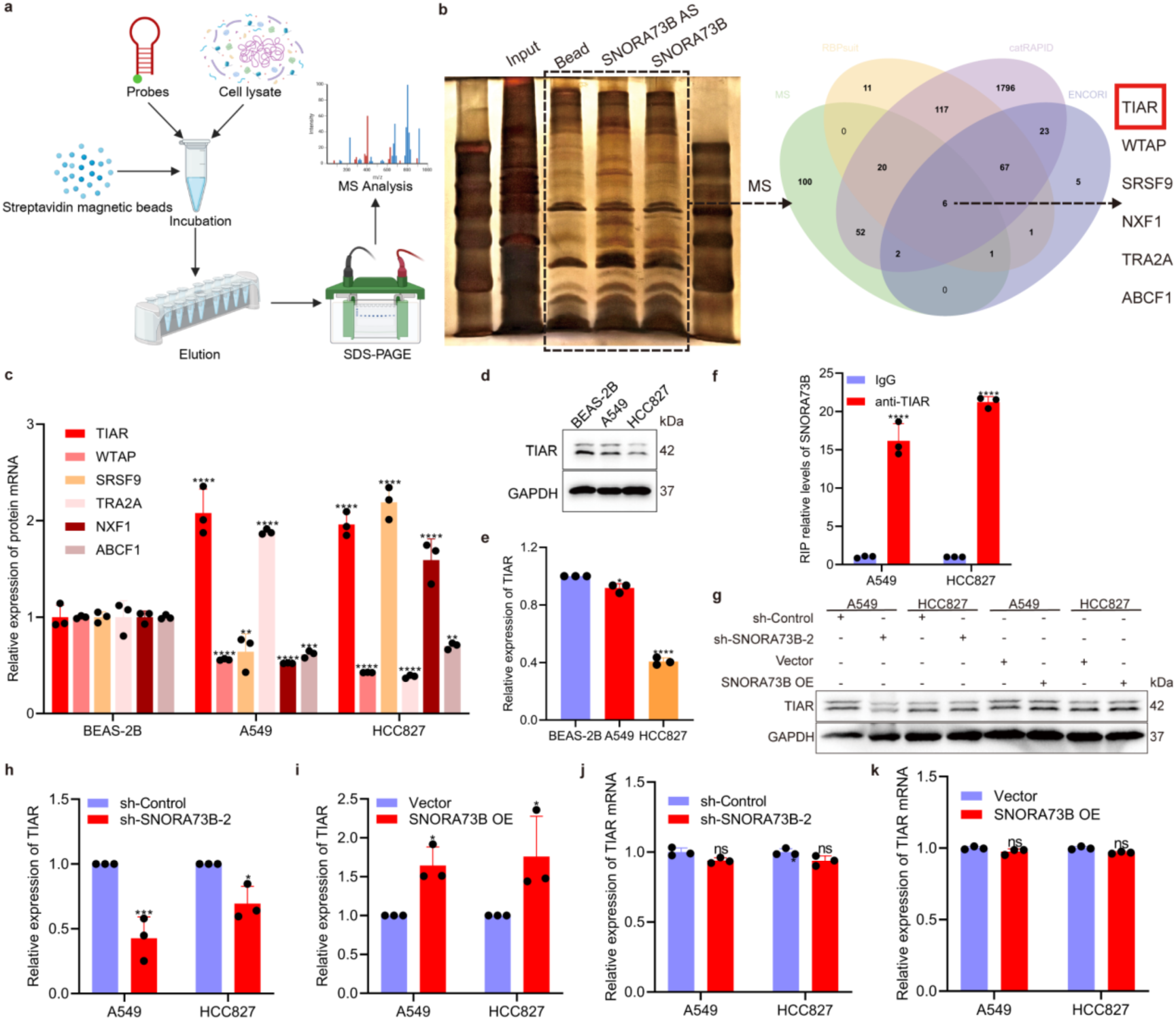
SNORA73B binds TIAR and upregulates its protein abundance. (**a**) Schematic of the biotinylated RNA pull-down and mass spectrometry workflow. (**b**) Silver staining SDS-PAGE gel of pull-down proteins (left) and Venn diagram of mass-spectrometry-identified candidates overlapping with RBPs from the RBPsuit, catRAPID, and ENCORI databases, yielding six shared hits (right). (**c**) RT-qPCR analysis of mRNA levels of the six candidate RBPs in BEAS-2B, A549, and HCC827 cells. (**d,e**) Western blot (**d**) and quantification (**e**) of TIAR protein in the three cell lines. (**f**) RIP-qPCR showing enrichment of SNORA73B by anti-TIAR antibody relative to IgG. (**g-i**) Western blot (**g**) and quantification of TIAR protein following SNORA73B knockdown (**h**) or overexpression (**i**) in A549 and HCC827 cells. (**j,k**) RT-qPCR analysis of TIAR mRNA after SNORA73B knockdown (**j**) or overexpression (**k**). Data are presented as the mean ± SD, n = 3 independent experiments. Statistical significance was determined by one-way ANOVA (**e**) or two-way ANOVA (**c,f,h,i,j,k**). Ns, not significant, \**P* < 0.05, \*\**P* < 0.01, \*\*\**P* < 0.001, \*\*\*\**P* < 0.0001.

To prioritize a functionally relevant binding partner for SNORA73B, we profiled basal mRNA expression of the six candidates in BEAS-2B, A549 and HCC827 cells. TIAR mRNA was highly expressed in both LUAD cell lines and was therefore selected for further study (**Fig. 6c**). Notably, TIAR protein abundance was lower in A549 and HCC827 cells than in BEAS-2B cells (**Fig. 6d,e**), despite elevated transcript levels, suggesting post-translational regulation or interaction with its binding partner of SNORA73B. RNA immunoprecipitation (RIP) with an anti-TIAR antibody significantly enriched SNORA73B compared with control IgG (**Fig. 6f**), confirming a direct interaction. Modulation of SNORA73B expression revealed that SNORA73B knockdown decreased TIAR protein, whereas SNORA73B overexpression significantly increased TIAR protein abundance in both cell lines (**Fig. 6g-i**). In contrast, TIAR mRNA levels remained unchanged upon SNORA73B perturbation (**Fig. 6j-k**). Thus, SNORA73B specifically binds TIAR and positively regulates its protein abundance without altering its mRNA, uncovering a post-transcriptional regulatory mechanism in LUAD.

### SNORA73B-TIAR-MYC signaling promotes malignant phenotypes via AKT phosphorylation

The BioGRID analysis of the TIAR interactome identified MYC as a candidate binding partner (**Fig. 7a**), consistent with evidence that TIAR binds the 3’ untranslated region (3’UTR) of MYC mRNA to regulate its transliation^32^. Given that c-Myc is a key node in the PI3K/AKT pathway^33^, we hypothesized that SNORA73B modulates c-Myc expression through TIAR to influence AKT phosphorylation. In A549 and HCC827 cells, SNORA73B knockdown decreased TIAR and c-Myc protein levels and reduced phosphorylated AKT (pAKT), without affecting total AKT (tAKT) (**Fig. 7b-f**). Conversely, SNORA73B overexpression significantly increased TIAR, c-Myc and pAKT (**Extended Data Fig. 5a-d**). These results indicate that SNORA73B upregulates c-Myc via TIAR, thereby enhancing AKT phosphorylation.

**Fig. 7.**
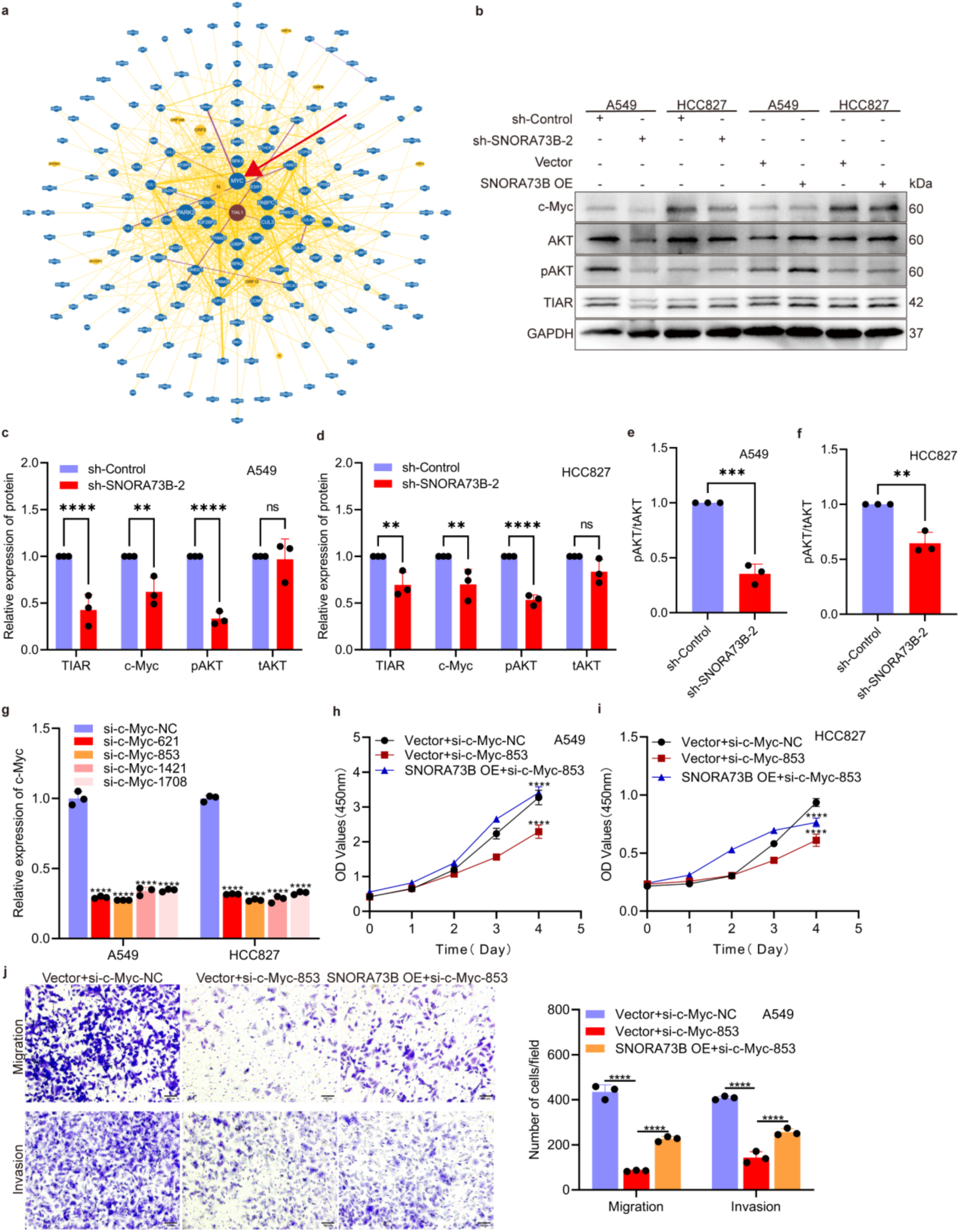
SNORA73B-TIAR--Myc signalling promotes malignant phenotypes via AKT phosphorylation. (**a**) Protein interaction network of TIAR (also known as TIAL1) from the BioGRID dataset; MYC is indicated as a candidate interacting protein. (**b-d**) Western blot and quantification of TIAR, c-Myc, pAKT and tAKT in A549 and HCC827 cells following SNORA73B knockdown or overexpression. (**e,f**) Quantification of pAKT/tAKT ratios in A549 (**e**) and HCC827 (**f**) cells upon SNORA73B knockdown. (**g**) RT-qPCR validation of c-Myc siRNA knockdown efficiency. (**h,i**) CCK-8 proliferation assays in A549 (**h**) and HCC827 (**i**) cells transfected with si-c-Myc alone or together with SNORA73B overexpression. (**j**) Transwell migration and invasion assays in A549 cells under the same conditions. Scale bar, 100 μm. Data are presented as the mean ± SD, n = 3 independent experiments (**b-g,j**) or n = 5 per group (**h,i**). Statistical significance was determined by two-way ANOVA (**b–d,g–j**) or two-tailed unpaired Student’s *t*-test (**e,f**). Ns, not significant, \*\**P* < 0.01, \*\*\**P* < 0.001, \*\*\*\**P* < 0.0001.

To determine whether c-Myc mediates SNORA73B-driven malignant cell phenotypes, we designed four siRNAs targeting *c-Myc*; si-c-Myc-853 achieved the greatest knockdown (**Fig. 7g**) and was used in rescue experiments. Co-transfection of si-c-Myc-853 with SNORA73B-overexpressing cells partially reversed SNORA73B-induced proliferation (**Fig. 7h,i** and **Extended Data Fig. 5e**) as well as migration and invasion (**Fig. 7j** and **Extended Data Fig. 5f**). Collectively, these findings demonstrate that the SNORA73B-TIAR-Myc axis promotes LUAD cell proliferation, migration and invasion by activating downstream AKT phosphorylation.

### SNORA73B promotes LUAD tumour growth by regulating c-Myc

To validate the role of the SNORA73B-TIAR-c-Myc axis *in vivo*, we established subcutaneous xenografts in nude mice (experimental scheme in **Fig. 8a**). SNORA73B overexpression significantly increased tumour volume and weight relative to controls; concomitant c-Myc knockdown partially reversed this pro-tumorigenic effect **(Fig. 8b-d**). Further RT-qPCR analysis of excised tumours confirmed corresponding changes in SNORA73B and c-Myc expression (**Fig. 8e-g**). These data demonstrate that SNORA73B overexpression promotes LUAD tumour growth, and that this effect is partially dependent on c-Myc, reinforcing the critical role of the SNORA73B-TIAR-c-Myc axis in LUAD progression (**Fig. 8h**).

**Fig. 8.**
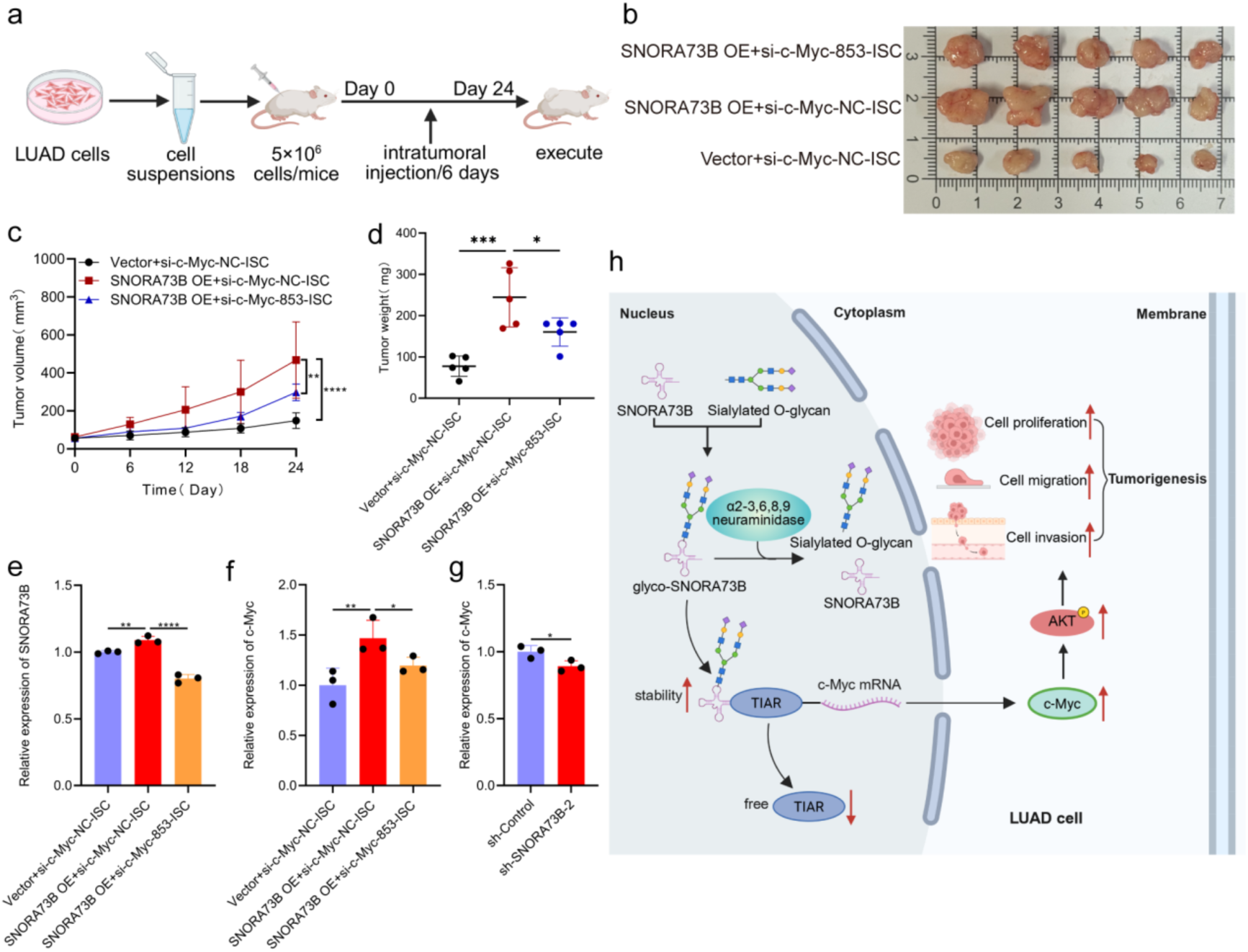
SNORA73B promotes LUAD tumour growth by regulating c-Myc. (**a**) Schematic of subcutaneous xenograft experiment. (**b**) Representative images of xenograft tumours from nude mice injected with Vector+si-c-Myc-NC-ISC, SNORA73B OE+si-c-Myc-NC-ISC, or SNORA73B OE+si-c-Myc-853-ISC. (**c**) Tumour volume over time in the indicated groups. (**d**) Tumour weight at endpoint. (**e,f**) RT-qPCR analysis of SNORA73 (**e**) and c-Myc (**f**) mRNA in xenograft tumours following SNORA73B overexpression and c-Myc knockdown. (**g**) RT-qPCR analysis of c-Myc mRNA in xenograft tumours after SNORA73B knockdown. (**h**) Schematic of the mechanism by which glyco-SNORA73B promotes LUAD progression via the activation of the TIAR/c-Myc/AKT signaling pathway. Data are presented as the mean ± SD, n = 3 independent experiments (**e-g**) or n = 5 mice per group (**b-d**). Statistical significance was determined by two-way ANOVA (**c,e,f**), one-way ANOVA (**d**) or two-tailed unpaired Student’s *t*-test (**g**). \**P* < 0.05, \*\**P* < 0.01, \*\*\**P* < 0.001, \*\*\*\**P* < 0.0001.

## Discussion

Since the discovery of cell-surface glycoRNAs, the field has progressed rapidly through methodological innovation. Early metabolic labeling confirmed that small ncRNAs bear N-glycans, but these approaches were limited by low labeling efficiency and lack of sequence resolution. Subsequent advances—including ARPLA for *in situ* imaging^21^, glycometabolic probe-based labelling^34^; data-independent acquisition (DIA)-based glycomic workflow (GlycanDIA)^35^, dual bioorthogonal labeling strategies^36^, and surface RNA amplification methods such as AMOUR and Intact-Surface-FISH^37^—have enabled sensitive detection and spatial mapping of glycoRNAs. Despite this progress, most studies remain limited to the population level, unable to resolve the sequence identity or function of individual glycoRNA species^38^. Although both N- and O-glycans have been detected on RNA^39^, their functional significance in cancer has remains speculative. By integrating metabolic labeling, glycomic mass spectrometry and RNA blotting, we directly anchor glycosylation to SNORA73B, a snoRNA with established oncogenic activity. We provide evidence that SNORA73B carries large O-glycans, making it one of the few ncRNAs with defined sequence identity and function. This study thus shifts glycoRNA research from population characterisation to molecular-level functional understanding, establishing a direct mechanistic link between a specific glycoRNA and tumour biology.

SnoRNAs have traditionally been viewed as guides for rRNA modification^40,41^, however, a growing number exhibit non-classical functions^42^, participating in tumorigenesis through protein interactions or gene expression. For instance, SNORA37^43^, SNORA38B^44^, SNORD88^45^ and SNORA49^46^ can all promote tumour progression via non-classical mechanisms. We find that SNORA73B promotes LUAD malignant progression not by directing rRNA modification but rather by engaging the TIAR-c-Myc-AKT signalling axis, consistent with this emerging functional paradigm. Importantly, we uncover an additional layer of regulation: glycosylation of SNORA73B likely modulates its stability or its ability to interact with TIAR. This discovery integrates two cutting-edge fields—RNA chemical modifications and non-canonical snoRNA biology—and introduce a “modification-dependent” dimension to snoRNA function. Consequently, evaluating snoRNAs as therapeutic targets requires consideration not only of their sequence but also of their modification status and their functional consequences.

RNA modifications are central to epigenetic regulation^47^; among them, small-molecule modifications such as *N*^6^-methyladenosine (m^6^A) have been extensively characterized^48,49^. By contrast, glycosylation involves the covalent attachment of structurally complex macromolecular glycans to RNA, endowing RNA modifications. Although recent studies have reported the regulation of RNA with glycoprotein-like molecular recognition properties. Recent work on redox-responsive RNA acylation^50^ demonstrate the feasibility of attaching functional groups to RNA, but naturally occurring glycosylation goes further. GlycoRNA can engage lectin and lectin-like receptor families through their glycan moieties, influencing processes including tumour proliferation and metastasis^51^, immune evasion^52^ and internalization of cell-penetrating peptides^53^. Our study extends this paradigm to cancer signalling: glycosylation on SNORA73B may govern its recognition by TIAR or other effector molecules, thereby controlling c-Myc-AKT pathway activation. This reveals a model of action that transcends classical epigenetic regulation—RNA molecules can acquire “lectin ligand” functionality through covalently linked glycosylated glycans, enabling them to participate directly in signal transduction and intercellular communication. These findings provide a new framework for non-classical RNA functions in malignancy.

Dysregulated snoRNA expression in lung cancer is well documented^54,55^, and snoRNAs are increasingly recognized as potential liquid biopsy biomarkers^56^. We introduce a further dimension: glycosylated snoRNAs. Glyco-SNORA73B is significantly elevated in plasma from LUAD patients and shows diagnostic value even in early stage disease. Current biomarkers such as circulating tumor DNA (ctDNA) and carcinoembryonic antigen (CEA) have limited sensitivity in early detection^57,58^, whereas tumour-derived glycoRNAs stably present in plasma may offer a complementary strategy^59^. Glycosylated snoRNA possesses a unique “dual recognition” feature—specific sequence combined with specific modifications—that could confer greater plasma stability and tumour specificity, a notion supported by existing data^23^. New detection methods such as RNA-optimized periodate oxidation and aldehyde labeling (rPAL) provide the technical means to capture low-abundance glycoRNAs^30^. Glyco-SNORA73B therefore demonstrates potential advantages in early detection and diagnostic specificity over conventional nucleic acid or protein biomarkers, warranting validation in prospective clinical cohorts.

Several limitations should be addressed in future work. The glycosylation sites on SNORA73B remain unmapped, and the cognate glycosyltransferases and upstream regulatory networks are unknown, limiting a complete understanding of modification control. The molecular basis of glyco-SNORA73B recognition by TIAR awaits structural elucidation to define interaction interfaces and key residues. Our findings rely on cell lines and xenograft models; genetically engineered mouse models are needed to examine pathway physiology and immune microenvironment interactions. Finally, clinical translation of glyco-SNORA73B into a clinically applicable biomarker or therapeutic target will require large-scale, multicenter prospective studies, as well as preclinical development of targeting strategies. Resolving these questions will deepen our understanding of RNA glycosylation in cancer and open new avenues for the precision diagnosis and therapy in LUAD.

In summary, we systematically characterize the glycosylation profile of SNORA73B and demonstrate that it functions as an oncogenic glycoRNA that drives LUAD malignant progression through the TIAR-c-Myc-AKT axis. Its marked upregulation in LUAD tissues and plasma supports its utility as an early diagnostic biomarker. These findings expand the functional repertoire of snoRNAs in cancer and establish glyco-SNORA73B as a novel molecular target with both diagnostic and therapeutic promise in LUAD.

## Methods

### Cell culture

The human LUAD cell lines A549, HCC827, NCI-H1299 and 95-D, and the normal bronchial epithelial cell line BEAS-2B, were obtained from BaiDi Biotechnology Co., Ltd. All cells were maintained at 37℃ in a humidified atmosphere of 5% CO_2_. A549, 95-D and BEAS-2B cells were cultured in DMEM (10-013-CVR, Corning), and HCC827 and NCI-H1299 cells in RPMI 1640 medium (10-040-CVR, Corning). Both basal media were supplemented with 10% fetal bovine serum (FBS, ST30-3302, PAN) and 1% penicillin-streptomycin-amphotericin B (C0224, Beyotime).

### Clinical samples

This study was approved by the Medical Research Ethics Committee of Ningbo University (approval No. NBU-2024-316), and written informed consent was obtained from all participants. Between 2021 and 2025, plasma samples were prospectively collected from 66 patients with LUAD and 27 healthy controls (HCs) at the First Affiliated Hospital of Ningbo University. Paired tumour and adjacent normal tissue specimens were obtained from an independent cohort of 76 LUAD patients at the same hospital. None of the patients had received prior chemotherapy or radiotherapy. Tissue specimens were immediately submerged in RNA protection reagent (SB-MR083, ShareBio) and stored at -80°C until processing.

### RNA extraction and purification

Total RNA was isolated from tissues and plasma using Trizol LS reagent (10296010, Invitrogen), and from cultured cells using Trizol reagent (15596018, Invitrogen), following the manufacturers’ protocols. Small and large RNAs were differentially precipitated with the Quick-RNA™ Microprep Kit (R1050, Zymo Research). For further purification, RNA samples were bound to a Zymo RNA clean and concentrator column (C1004-50, Zymo Research) by mixing with 2 volumes of RNA Binding Buffer (R1013-2-25, Zymo Research) and 3 volumes of 100% ethanol, washed three times (the final two washes with 3 volumes of 100% ethanol), and eluted in nuclease-free water. RNA concentration and quality were measured on a DeNovix DS-11 spectrophotometer (DeNovix, DE), and samples were stored at −80°C.

### Click chemistry, RNA electrophoresis, blotting and imaging

Cells were metabolically labelled by culturing in medium containing 100 μM Ac_4_ManNAz (T875048, Macklin) for 36 h, with fresh compound added at 0 h and 36h. Total RNA was extracted, and glycosylated RNAs were conjugated to DBCO-PEG4-Biotin (D916831, Macklin) by copper-free strain-promoted azide-alkyne cycloaddition (SPAAC). For the SPAAC reaction, RNA samples were mixed with biotin and. Each reaction contained 9 μL RNA, 1 μL 10 mM DBCO-biotin and 10 μL dye-free gel loading buffer II (df-GLBII: 95% formamide, 18 mM EDTA, 0.025% SDS). After denaturation at 55°C for 10 min, the reaction was quenched with 80 μL water, followed by 2 volumes (200 μL) RNA Binding Buffer and 3 volumes (300 μL) 100% ethanol. The mixture was purified on a Zymo column as described above and analyzed by gel electrophoresis.

The purified biotin-labeled RNA was resuspended in 15 μL df-GLBII containing 1× SybrGold (S11494, Thermo Fisher Scientific), denatured again (55°C for 10 min, and then on ice for 3 min), and resolved on a 1% agarose formaldehyde denaturing gel in 1× MOPS buffer at 110 V for 45 min. Total RNA was visualized with a gel imager (Bio-Rad) and transferred to a 0.45-μm nitrocellulose membrane (10600002, Cytiva Amesham) at room temperature for 2 h. The membrane was crosslinked with UV-C light, blocked with Odyssey Blocking Buffer (927-70001, Li-Cor Biosciences) for 45 min, incubated with streptavidin-IR800 (926-32230, Li-Cor Biosciences; 1:10,000 in blocking buffer) for 30 min, washed three times with PBS containing 0.1% Tween-20 (0777-1L, Amresco) for 5 min each, briefly rinsed with PBS, and imaged with an Odyssey CLx scanner (Li-Cor Biosciences).

For RNA blot detection of SNORA73B, a biotin-labeled probe (**Supplementary Table 2**) was synthesized by Ningbo Laiqiao Biotechnology Co., Ltd, and the Biotin Northern Blot Kit (R0219, Beyotime) was used following the manufacturer’s protocol.

### Small RNA enrichment and microarray analysis

Small RNAs were extracted and purified from Ac_4_ManNAz-labelled A549 or BEAS-2B cells. An aliquot of 500 ng was retained as the input control, and 25 μg was biotinylated by SPAAC as described above. For enrichment, MyOne C1 Streptavidin beads (65001, Thermo Fisher Scientific) were blocked for 1 h at 25°C in biotin wash buffer (10 mM Tris-HCl pH 7.5, 1 mM EDTA, 100 mM NaCl, 0.05% Tween-20) supplemented with 50 ng/μL glycogen (AM9510, Thermo Fisher Scientific). The biotinylated RNA was diluted to 33 ng/μL in 750 μL biotin wash buffer and incubated with blocked beads at 4°C for 2 h. Beads were then washed twice with 1 mL ChIRP wash buffer (2x SSC, 0.5% SDS), twice with 1 mL biotin wash buffer, and twice with 1 mL NT2 buffer (50 mM Tris-HCl pH 7.5, 150 mM NaCl, 1 mM MgCl_2_, 0.005% NP-40), for 3 min each at 25°C. The steptavidin-bound RNA and the input RNA were subjected to small RNA microarray analysis (Human Small RNA Array V1.0, Aksomics).

### Bioinformatics analyses

The differentially expressed RNAs in LUAD were analysed using the SangerBox platform (http://sangerbox.com/)^60^. Briefly, TCGA LUAD transcriptome data were retrieved, and the limma R package (version 3.40.6) was used to compare tumour versus adjacent normal tissues. From the small RNA microarray data, glycosylated snoRNAs that were consistently upregulated in both A549 and BEAS-2B cells were selected using thresholds of |FC| ≥ 2 and *P* < 0.05. Visualization was generated on the same platform. The secondary structure of human SNORA73B was predicted with RNAcentral (https://rnacentral.org/).

### RT-qPCR

Complementary DNA (cDNA) was synthesized using the NovoScript® Plus All-in-one 1st Strand cDNA Synthesis SuperMix (gDNA Purge) (E047-01B, Novoprotein). Quantitative polymerase chain reaction (qPCR) was performed with the NovoScript® SYBR qPCR SuperMix Plus Kit (E096-01A, Novoprotein) on a Roche LightCycler® 480 II real-time system (Roche Diagnostics) following the manufacturer’s instructions. Primer sequences are provided in **Supplementary Table 3**.

### ROC analysis

ROC analysis was performed to evaluate the diagnostic performance of SNORA73B. The Yorden index (J = sensitivity + specificity - 1) was used to determine the optimal cut-off value. Diagnostic accuracy was quantified by the AUC.

### SnoRNA localization assay

Nuclear and cytoplasmic RNA were isolated using the Paris Kit (AM1921, Thermo Fisher Scientific) according to the manufacturer’s instructions. RT-qPCR was performed to quantify U6, GAPDH, and SNORA73B levels in nuclear and cytoplasmic fractions. Primer sequences are provided in **Supplementary Table 3**.

### Cell transfection

The pLV3-CMV-MCS2-EF1a-CopGFP-Puro (Vector) and pLV3-CMV-SNORA73B (human)-snoRNA-EF1a-CopGFP-Puro (SNORA73B OE) plasmids were obtained from MiaoLingPlasmid. The plko.1-copGFP-PURO (sh-Control), plko.1-copGFP-PURO-sh-SNORA73B-1, -2, and -3 (sh-SNORA73B-1, -2, -3) from Beijing Tsingke Biotech Co., Ltd. siRNAs targeting c-Myc (si-c-Myc-NC, -621, -853, -1421, -1708) were obtained from Shanghai GenePharma Co., Ltd. To generate stably expressing cell lines, LUAD cells were transfected with Lipofectamine® 2000 Reagent (11668-019, Invitrogen) and selected with puromycin (P8230, Solarbio).

### GlycoRNA glycomics mass spectrometry

Total RNA was isolated from A549 cells expressing sh-Control and sh-SNORA73B-2 and subjected to glycomics mass spectrometry (Guangzhou Epibiotek). Relative quantification of O- and N-linked glycans were compared between the two groups using a two-tailed unpaired Student’s *t*-test, glucan structures with *P* < 0.05 were considered differentially expressed (**Supplementary Tables 4,5**).

### Sialic acid removal

For sialic acid removal, total RNA (20 μg) was incubated with 2 μL ofα2-3,6,8,9 neuraminidase (50 U/mL, P0722S, NEB) and 1 μL of GlycoBuffer 1 at 37°C for 90 min.

### CCK-8 assay

Following transfection, cells were trypsinized, counted and seeded into 96-well plates at 2 × 10³ cells per well in 100 μL of culture medium. After 4 h of adhesion (day 0), 10 μL of CCK-8 solution (GK10001, Glpbio) was added to the wells at day 0, 1, 2, 3 and 4. The plate was incubated for 2 h at 37°C in the dark, and the absorbance at 450 nm was recorded using a microplate reader (Thermo Fisher Scientific).

### Colony formation assay

Transfected cells were trypsinized, counted and seeded at 1,500 cells per well in 6-well plates. Cells were cultured for 14 days with medium replacement once per week. Subsequently, cells were washed twice with PBS, fixed in 4% paraformaldehyde (P1110, Solabio) for 30 min, and stained with 0.1% crystal violet (G1063, Solabio) for 30 min. After rinsing with water, plates were dried, photographed, and colonies were counted.

### Transwell migration and invasion assays

For both assays, 600 μL of culture medium supplemented with 20% FBS was added to the lower chamber (3421, Corning). Transfected cells were trypsinized, counted, and 3 × 10⁴ cells were resuspended in serum-free medium and seeded into the upper chamber. For invasion assays, the upper chamber was pre-coated with Matrigel (356234, Corning) diluted to 250 μg/mL in serum-free medium and incubated at 37°C for at least 1 h to solidify before cell seeding. Migration assays were cultured for 24 h and invasion assays for 48 h. After incubation, non-migrated cells were removed from the upper chamber surface, and cells that had migrated or invaded were stained with 0.1% crystal violet, air-dried and counted under a light microscope.

### Biotin-labelled RNA pull-down and mass spectrometry

Biotin-labeled sense and antisense SNORA73B probes (**Supplementary Table 6**) were synthesized (Shanghai GenePharma) and used to pull down SNORA73B-binding proteins with the Biotin-labelled RNA Pull-Down Kit (FI8702, Fitgene). Protein complexes were resolved by SDS-PAGE (PG112, Epizyme), and visualized by silver staining (PAGE Gel Silver Staining Kit, G7210, Solarbio). Bands of interest were excised and analyzed by mass spectrometry (Shanghai iProteome Biotechnology).

### RNA immunoprecipitation (RIP)

RIP was performed using the Magna RIP™ RNA-Binding Protein Immunoprecipitation Kit (17-700, Millipore) following the manufacturer’s instructions. Briefly, cell lysates were incubated overnight at 4°C with magnetic beads and anti-TIAR antibody (1:100, 8509, Cell Signaling Technology). After washing, co-precipitated RNA was purified and analyzed by RT-qPCR.

### Western blotting

Cells were cultured in 6-well plates to 80-90% confluence and lysed in RIPA buffer (R0010, Solarbio). Protein concentration was determined with a BCA Protein Assay Kit (WB6501, NCM Biotech). Protein samples (30 μg) were desaturated at 100°C for 5 min, separated on 10% SDS-PAGE gels (80 V for 15 min, then 120 V for 60 min) and transferred to PVDF membranes by wet transfer (Bio-Rad) for 2 h. Membranes were blocked with 5% non-fat milk for 3 h at room temperature and then incubated overnight at 4°C with primary antibodies: anti-TIAR (1:1000, 8509, Cell Signaling Technology, Inc), anti-c-Myc (1:7,000, 10828-1-AP, Proteintech), anti-tAKT (1:1,000, sc-81434, Santa Cruz Biotechnology), anti-pAKT (1:1,000, sc-514032, Santa Cruz Biotech), and anti-GAPDH (1:1,000, sc-47724, Santa Cruz Biotech), respectively. After three 10-min washes with TBST, membranes were incubated with HRP-conjugated secondary antibodies (1:6,000, RGAR001 or RGAM001, Proteintech) for 2 h at room temperature, followed by three additional TBST washes. Signals were detected with an Enhanced Chemiluminescent Reagent Kit (P10100, NCM Biotech), and bands intensities were quantified using ImageJ.

### Xenografts in nude mice

All animal procedures were approved by the Animal Ethics Committee of Ningbo University (approval No. NBU-2024-12922). Twenty-five male BALB/c nude mice (3 weeks old; GemPharmatech) were randomly assigned to five groups (n = 5 per group) and housed in the Ningbo University Laboratory Animal Center. A549 cells stably expressing sh-Control, sh-SNORA73B-2, empty vector (Vector), or SNORA73B overexpression (SNORA73B OE) were injected subcutaneously into the axillary region (5 × 10⁶ cells per mouse). The five groups were: (1) sh-Control; (2) sh-SNORA73B-2; (3) Vector + si-c-Myc-NC-ISC (negative control siRNA); (4) SNORA73B OE + si-c-Myc-NC-ISC; and (5) SNORA73B OE + si-c-Myc-853-ISC. For groups 3-5, intratumoral injections of indicated siRNA were administered every 6 days. Tumour dimensions (longest diameter *L* and shortest diameter *W*) were measured with a caliper, and tumour volume was calculated as (*L* × *W*²) /2. After 24 days, mice were euthanized, tumours were excised, photographed, and analyzed.

### Statistical analysis

Data were analyzed using GraphPad Prism 9 and are presented as mean ± SD. The statistical tests used are specified in the figure legends. *P* < 0.05 was considered statistically significant.

## Availability of data and material

The small RNA microarray data generated in this study have been deposited in the Open Archive for Miscellaneous Data (OMIX) of the National Genomics Data Center (NGDC) under accession code OMIX017136 (https://ngdc.cncb.ac.cn/omix/preview/1xz4wYqi). All other data supporting the findings of this study are available from the corresponding author on reasonable request.

## Acknowledgements

We thank all members of the Z.G. laboratory for valuable discussion. We acknowledge the staff from the Laboratory Animal Center and Large-scale Instrument Platform at Ningbo University for their technical support.

## CRediT authorship contribution statement

**Lulu Yang**: Conceptualization, Data curation, Formal analysis, Investigation, Methodology, Validation, Writing—original draft. **Boyang Wang**: Investigation, Methodology, Validation. **Yufei Sheng, Zhuobin Deng, Jiali Liu, Zhiqi Hong and Lei Zheng**: Investigation, Methodology. **Chengwei Zhou and Wentao Hu**: Resources, Writing—review & editing. **Zhaohui Gong**: Conceptualization, Data curation, Supervision, Funding acquisition, Project administration, Writing—review and editing.

## Extended data

**Extended Data Fig. 1.**
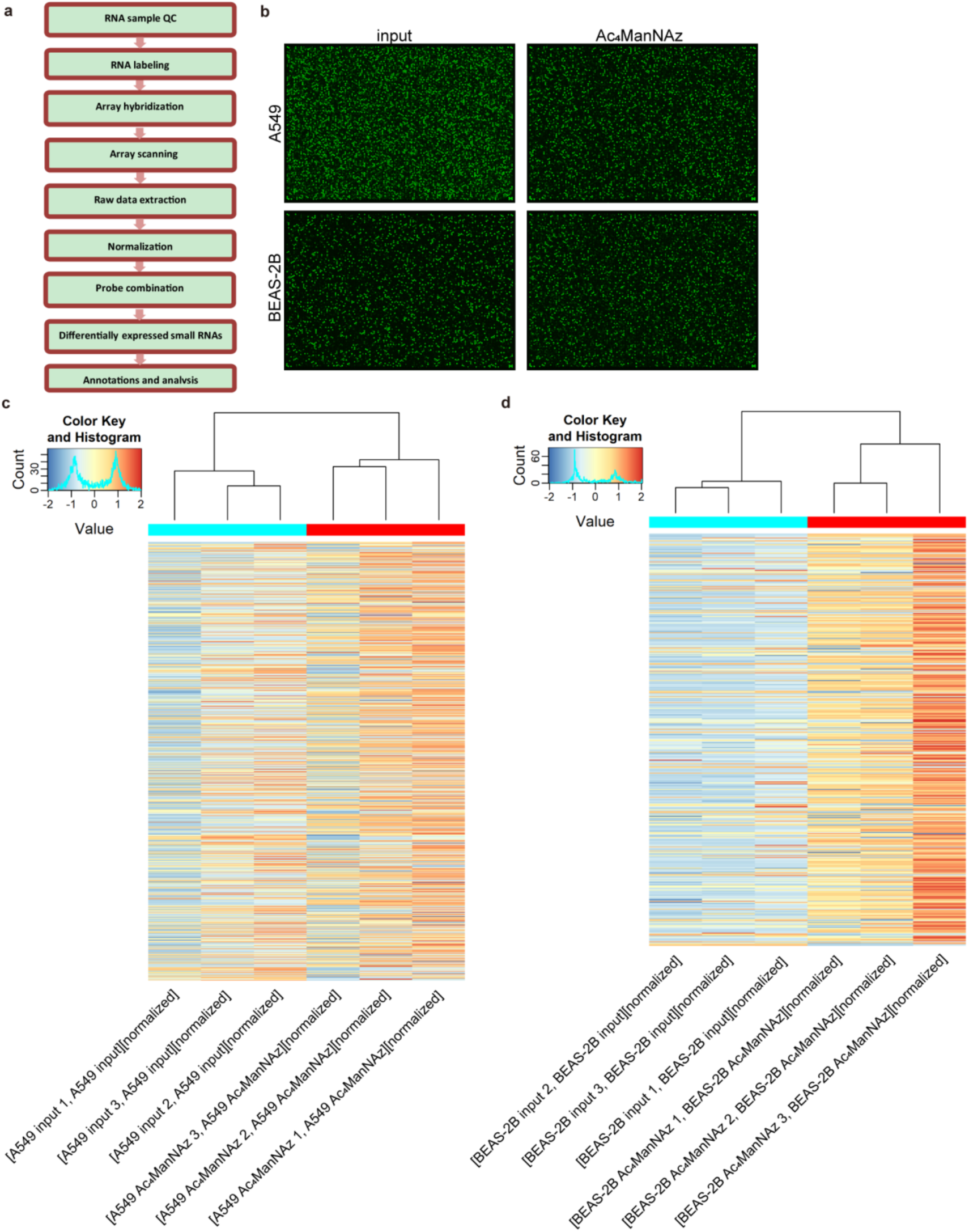
Small RNA microarray analysis reveals that snoRNA is a component of glycoRNA. (**a**) Schematic diagram of the workflow for small RNA microarray hybridization, signal detection, and data analysis. (**b**) Representative array images of small RNA (input group) and glycoRNA enriched by streptavidin pull-down (Ac4ManNAz group) from A549 and BEAS-2B cells following small RNA microarray hybridization. (**c,d**) Hierarchical clustering heatmap analysis of snoRNAs in A549 cells (**c**) and BEAS-2B cells (**d**). Each row in the figure represented a snoRNA, and each column represented a sample; the relative expression levels of snoRNAs were indicated by a color gradient (red: high expression; blue: low expression, see the legend in the upper left). The dendrogram at the top showed the clustering relationships based on the similarity of expression profiles among samples, and the color bar at the top indicated the experimental group to which each sample belonged (input group or Ac4ManNAz group).

**Extended Data Fig. 2.**
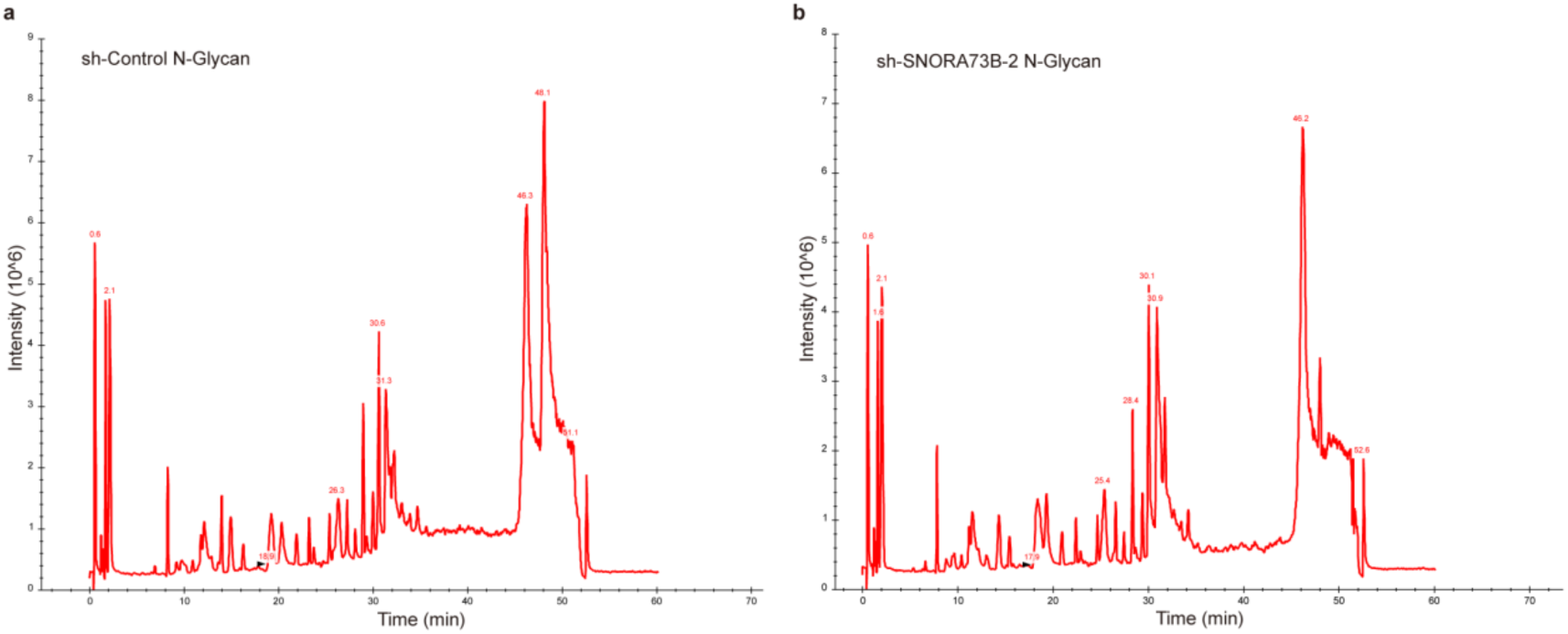
Glycomics MS analysis reveals that SNORA73B carries N-glycans. (**a**) Total ion chromatogram (TIC) of N-glycans in samples from the control group (sh-Control). (**b**) TIC of N-glycans in samples from the SNORA73B knockdown group (sh-SNORA73B-2).

**Extended Data Fig. 3.**
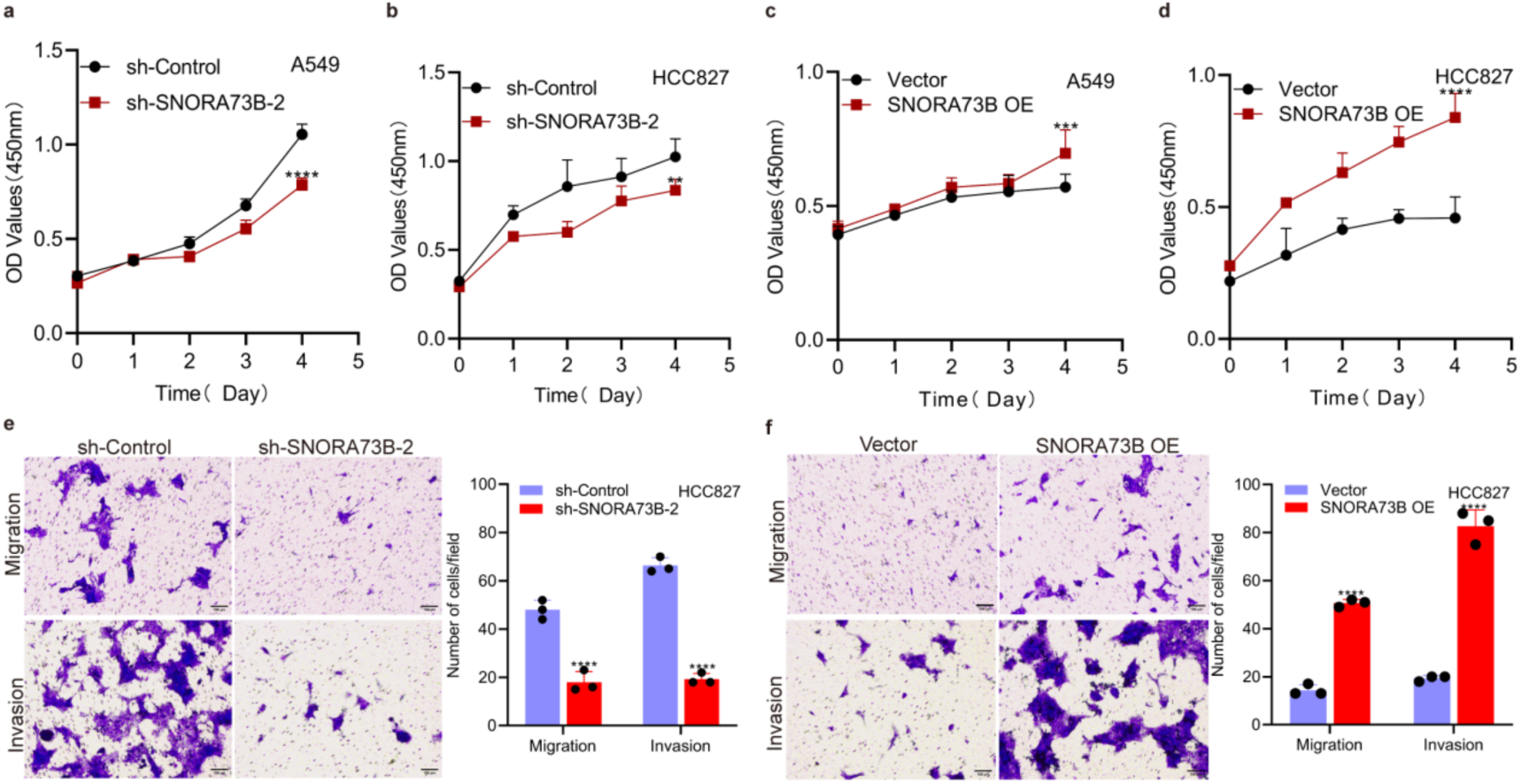
SNORA73B promotes the proliferation, migration and invasion of LUAD cells. (**a,b**) CCK-8 assays in A549 (**a**) and HCC827 (**b**) cells transfected with SNORA73B knockdown. (**c,d**) CCK-8 assays in A549 (**c**) and HCC827 (**d**) cells transfected with SNORA73B overexpression. (**e,f**) Transwell migration and invasion assays of HCC827 cells with SNORA73B knockdown (**e**) or overexpression (**f**). Scale bar, 100 μm. Data are presented as the mean ± SD, n = 5 per group (**a-d**) or n = 3 independent experiments (**e,f**). Statistical significance was determined by two-way ANOVA. \*\**P* < 0.01, \*\*\**P* < 0.001, \*\*\*\**P* < 0.0001.

**Extended Data Fig. 4.**
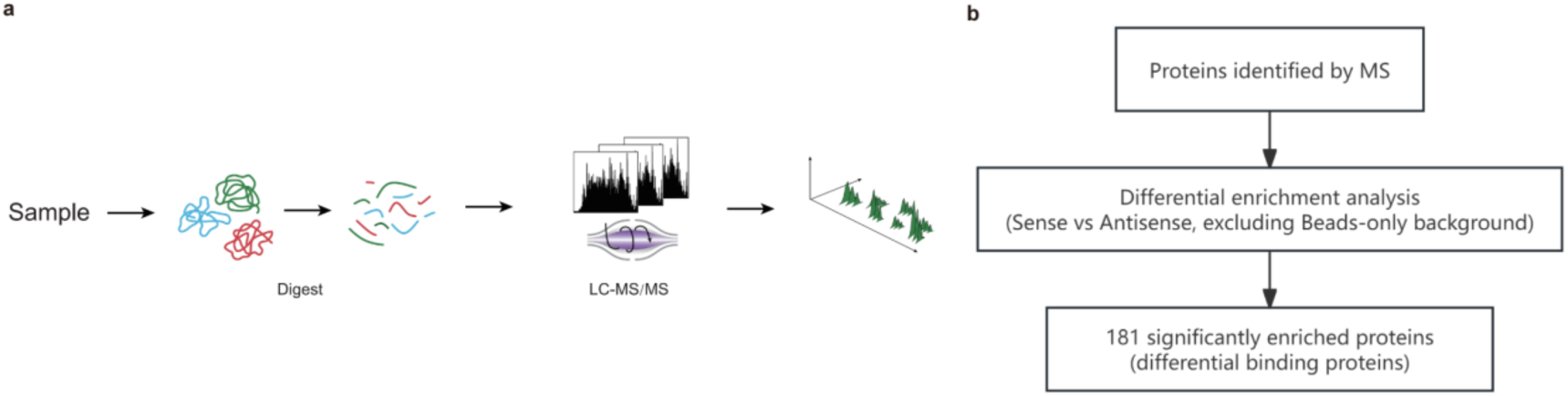
Identification of SNORA73B-interacting proteins using RNA pull-down combined with MS. (**a**) Flowchart of the label-free quantitative proteomics detection. (**b**) Flowchart for the MS-based screening of differentially binding proteins of SNORA73B.

**Extended Data Fig. 5.**
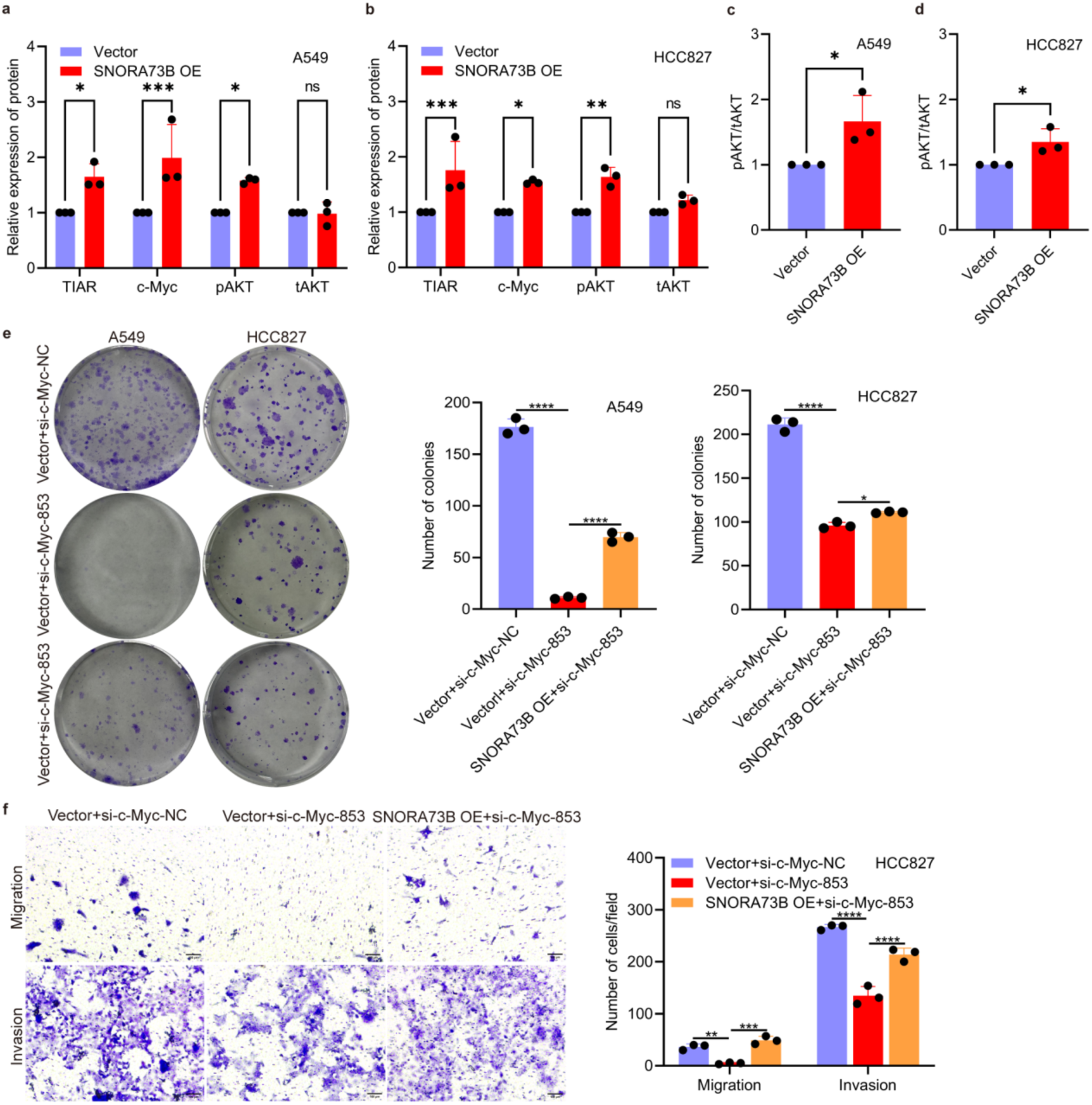
SNORA73B regulates AKT phosphorylation via the TIAR/c-Myc axis, thereby influencing the malignant cellular phenotype. (**a,b**) Western blot and quantification of TIAR, c-Myc, pAKT and tAKT in A549 (**a**) and HCC827 (**b**) cells following SNORA73B overexpression. (**c,d**) Quantification of pAKT/tAKT ratios in A549 (**c**) and HCC827 (**d**) cells upon SNORA73B overexpression. (**e**) Colony formation assays in A549 and HCC827 cells transfected with si-c-Myc alone or together with SNORA73B overexpression. (**f**) Transwell migration and invasion assays in HCC827 cells under the same conditions. Scale bar, 100 μm. Data are presented as the mean ± SD, n = 3 independent experiments. Statistical significance was determined by two-way ANOVA (**a,b,f**), one-way ANOVA (**e**) or two-tailed unpaired Student’s *t*-test (**c,d**). Ns, not significant, \**P* < 0.05, \*\**P* < 0.01, \*\*\**P* < 0.001, \*\*\*\**P* < 0.0001.

